# Gap-free nuclear and mitochondrial genomes of *Fusarium verticillioides* strain HN2

**DOI:** 10.1101/2022.12.22.521711

**Authors:** Wei Yang, Haoyu Zhai, Lei Yang, Qun Yang, Le Song, Jianyu Wu, Zhibing Lai, Guotian Li

## Abstract

Fusarium ear rot (FER) and Fusarium stalk rot (FSR) caused by the filamentous fungus *Fusarium verticillioides* have become increasingly serious around the world. Additionally, fumonisins produced by *F. verticillioides* threaten food and feed security. By adding the contribution of genomic resources to better understand the pathosystem including the mechanisms of *F. verticillioides*–maize interactions, and further improving the quality of the *F. verticillioides* genome, the gap-free nuclear genome and mitochondrial genome of *F. verticillioides* strain HN2 were sequenced and assembled. Using Oxford Nanopore long reads and next-generation sequencing short reads, the final 42.81-Mb genome was assembled into 12 contigs (N50 = 4.16-Mb). A total of 13,466 protein-coding genes were annotated, including 1,076 secreted proteins that contain 342 candidate effectors. In addition, we assembled the complete 53,764 bp mitochondrial genome. *F. verticillioides* strain 7600 genome assemblies are fragmented and high-quality reference genomes were needed. The genomes presented here will serve as an important resource for *F. verticillioides* research.

## Genome Announcement

Fusarium ear rot (FER) and Fusarium stalk rot (FSR) caused by the filamentous fungus *Fusarium verticillioides* pose serious threats to a wide range of crops, especially to maize, which lead to yield losses up to 30% (Li et al., 2019; Chen et al., 2021). In addition to crop losses, a number of notorious mycotoxins are produced by *F. verticillioides*. In addition to crop losses, upon infection, *F. verticillioides* produces notorious mycotoxins harmful to humans and animals (Bennett and Klich, 2003; Woloshuk and Shim, 2013). *F. verticillioides* infects maize at different growth stages (Blacutt et al., 2018), and volatile emissions from *F. verticillioides* attract insects to promote fungal infection and dispersal, making FER and FSR challenging to control (Franco et al., 2021). In maize, many studies on resistance to FER and FSR are focused on quantitative trait loci (QTL) (Samayoa et al., 2019; Wu et al., 2020). However, owing to the complex genetics of maize and the impact of different environmental conditions, resistance from QTL is unstable (Ju et al., 2017; Wen et al., 2021). Biological control for *F. verticillioides* is becoming a recent trend (Deepa et al., 2021). High-quality genomic resources provide valuable information on the pathogenesis of plant pathogens, which are crucial to reducing plant disease hazards and toxin contaminants. In 2010, the first genome of *F. verticillioides* strain 7600 was published (Ma et al., 2010), with a contigs number of 211. Given the fragmented assembly of the reference genome of strain 7600 as well as the need for a high-quality genome of *F. verticillioides* for the study of the *F. verticillioides*-maize pathosystem, we sequenced and assembled the gap-free nuclear and mitochondrial genomes of the *F. verticillioides* strain HN2, which contributes to the analysis of its characteristics in taxonomy, population genetics, basic and applied science.

The colony morphology of *F. verticillioides* strain HN2-isolated from a maize field at Zhengzhou, Henan Province, China-matched the published descriptions in 2005 (Supplementary Fig. S1) (Deepa and Sreenivasa, 2017). Using the cetyltrimethylammonium bromide method (Damm et al., 2008), we extracted high-quality DNA of 7-days-old strain HN2 grown on potato dextrose agar (PDA) plates under ambient light and at 28°C. We sequenced the internal transcribed spacer (ITS) region (Utami and Hiruma, 2022). Using the basic local alignment search tool (BLAST) with the NCBI database, we observed that strain HN2 is highly similar (99.8%) to that of *F. verticillioides* (Accession No. ON003540.1). We assembled a complete genome using a combination method of long reads and short reads. So, we sequenced the fungal genome with long and Illumina short reads using the Oxford Nanopore technology and the next-generation sequencing (NGS) technique (Wang et al., 2021; Zhao et al., 2021) at the Beijing Novogene Bioinformatics Technology Co., Ltd. The Oxford Nanopore long reads were obtained with the EXP-NBD104 kit and SQK-LSK109 connection kit of Oxford Nanopore Technologies Company from the PromethION platform following the manufacturer’s recommendations. NGS short reads were obtained with NEBNext® Ultra™ DNA Library Prep Kit for Illumina (NEB, USA) from the NovaSeq 6000 platform. To ensure the use of high-quality data, Nanopore long reads were filtered by removing reads with an average quality score less than 7 using Nanofilt (v2.6.0) (De Coster et al., 2018). Similarly, we filtered out the low-quality short reads that q value less than ‘20’ using fastp (v0.23.1) (Chen et al., 2018). Finally, we obtained 4.55-Gb clean Nanopore long reads and 2.46-Gb high-quality NGS short reads of *F. verticillioides* strain HN2. Before assembling, the genome size of strain HN2 was estimated to be 43.22-Mb using the *k*-mer method according to NGS short reads mapping with Jellyfish (v2.3.0) and GenomeScope (v2.0) (Supplementary Fig. S2) (Marcais and Kingsford, 2011; Ranallo-Benavidez et al., 2020). We assembled the nuclear genome using NextDenovo (v2.5.0) with these filtered reads. The draft genome base errors were further fixed using NextPolish (v1.4.0) (Hu et al., 2020). The final 42.81-Mb genome was assembled into 12 contigs (N50 = 4.16-Mb, GC = 47.89 %), which covers 99% of the estimated genome. Using a dual-locus phylogeny approach for phylogenetic assessment, including the translation elongation factor 1-α and β-tubulin gene regions (extracted from the assembled genomes), we identified strain HN2 as *F. verticillioides* within the *Fusarium fujikuroi* species complex by maximum parsimony analysis using PAUP (v4.0) with a heuristic search (Supplementary Fig. S3) (Swofford, 2003; Herron et al., 2015; Gardiner, 2018). Compared with the reference genome of *F. verticillioides* strain 7600 (211 contigs, N50 = 0.39-Mb) (Table 1), the genome assembly showed a notable improvement (Ma et al., 2010). The RaGOO (v1.1) scaffolding approach was used to anchor these contigs into 12 scaffolds, including 11 gap-free chromosomes and the supernumerary scaffold (super_scaffold) (Blacutt et al., 2018; Alonge et al., 2019). Using D-GENIES (Cabanettes and Klopp, 2018), the 11 chromosomes of strain HN2 show high linear relationships with the reference genome of *F. verticillioides* strain 7600, except for the inversions on chromosomes 3, 10 and 11 (Supplementary Fig. S4) (Ma et al., 2010). Among the 11 chromosomes, telomeric satellite sequences (TTAGGG/CCCTAA) were identified for both ends of chromosomes 7, and 11 and for one end of chromosomes 1, 2, 3, 4, 6, 9, and 10 (Supplementary Table S1). In addition, the super_scaffold (721,913 bp) with telomeric satellite sequence at both ends is presumed to be a conditionally dispensable chromosome. RepeatMasker (v4.1.2) (Tarailo-Graovac and Chen, 2009) and RepeatModeler (v2.0.3) (Flynn et al., 2020) were used to identify and mask 1.52-Mb (3.56 %) repetitive sequences in the genome (Supplementary Table S2). The genome assembly is at the high-quality level based on the BUSCO (v5.3.2) score at 99.2% (Supplementary Table S3) (Simao et al., 2015). We aligned the short NGS reads to the genome of strain HN2 and obtained a mapping rate of 97.38%, which further indicates its high-quality genome assembly, visualized using TBtools (v1.098761) (Chen et al., 2020).

**Table 1.** Summary of the *Fusarium verticillioides* strain HN2 genome

| Characteristics<sup>a</sup> | Strain |  |
| --- | --- | --- |
|  | HN2 | 7600 |
| Sequencing technology | Nanopore,<br>Illumina | Sanger |
| Genome assembly size (bp) | 42,814,391 | 41,844,914 |
| Contig number | 12 | 211 |
| Contig N50 (bp) | 4,158,290 | 392,397 |
| GC content (%) | 47.89 | 48.69 |
| Predicted genes | 13,466 | 14,179 |
| Secreted proteins | 1,076 | 1,549 |
| BUSCO completeness (%) | 99.2 | 98.5 |

<sup>a</sup> BUSCO, benchmarking universal single-copy orthologs.

We used 3.70-Gb RNA-seq data of *F. verticillioides* strain 7600 at the NCBI database (Ma et al., 2010) to fully annotate the genome. A total of 13,466 genes were predicted to be protein-coding genes using Funannotate (v1.8.9) which includes *ab initio* gene predictor (Augustus and GeneMark for predictions), and among them 13,055 (97%) were functionally annotated with the eggNOG database (Huerta-Cepas et al., 2019). We identified a total of 10,531 one-to-one orthologs between the genome of strain 7600 and HN2 using OrthoFinder (v2.5.2) (Emms and Kelly, 2019). Using tRNAscan-SE (v2.0.9) tool for transfer RNAs (tRNAs) detection (Chan and Lowe, 2019), we identified 297 tRNAs with the average length of 88 bp. Meanwhile we identified 66 ribosomal RNAs (rRNAs) using RNAmmer (v1.2) (Lagesen et al., 2007). In total, 28 small nuclear RNAs (snRNAs), 3 small RNAs (sRNAs) were identified using Infernal (v1.1.4) based on the Rfam database with default parameters (Nawrocki et al., 2009). The clean short reads of strain HN2 were mapped to the reference genome of strain 7600 with a mapping rate of 95.74% (Ma et al., 2010). We performed single nucleotide polymorphism (SNPs) and Insertion/Deletion (InDels) calling using Picard (v2.27.1) (https://broadinstitute.github.io/picard/) and GATK (v4.2.6.1) (McKenna et al., 2010). We used Lumpy (v0.2.13) to identify large structural variants (SVs) between strains HN2 and 7600 (Ma et al., 2010; Layer et al., 2014). Subsequently, we used the ANNOVAR tool to annotate SNPs, InDels, and SVs (Supplementary Table S4) (Wang et al., 2010). By performing the pipeline above, we obtained 241,434 SNPs, 24,577 InDels, and 3,793 SVs. We identified 1,076 secreted proteins using SignalP (v5.0) (Armenteros et al., 2019) and TMHMM (v2.0). Most secreted proteins are concentrated towards both ends of each chromosome, except for 218 secreted proteins on chromosomes 10 and 11. Using EffectorP (v3.0), we predicted 342 candidate effectors (Sperschneider and Dodds, 2022), including 98 cytoplasmic effectors and 244 apoplastic effectors. We analyzed whether the 342 candidate effectors of *F. verticillioides* strain HN2 are conserved among fungal pathogens of maize (Supplementary Table S5). The results showed that 332 candidate effectors were identified in *F. verticillioides* strains 7600 and BRIP14953, 272 in *Fusarium graminearum* strain PH-1, 134 in *Bipolaris maydis* strain C5, and 131 in *Exserohilum turcicum* strain Et28A (Ma et al., 2010; Ohm et al., 2012; Condon et al., 2013; King et al., 2015; Gardiner, 2018). Thus, a total of 122 candidate effectors were conserved in five maize fungal pathogens (Fig. 1). Analyzing the 342 candidate effectors of *F. verticillioides* strain HN2 with the Pathogen-Host Interaction (PHI) database with the parameter ‘-evalue 1.0e-5’ (Urban et al., 2022), we obtained 116 matched proteins, some of which are involved in fungal virulence (Rogers et al., 2000; Lu and Faris, 2019). Secondary metabolism gene clusters play an important role in toxin biosynthesis (Osbourn, 2010). We used the fungal version of antiSMASH (Blin et al., 2021), and predicted 54 secondary metabolism gene clusters, including 16 polyketide synthases (PKS), 12 non-ribosomal peptide synthetases (NRPS), 12 NRPS-like fragments (NPRS-Like), 3 indole synthases, and 11 terpene synthases. Carbohydrate-active enzymes (CAZymes) are key for fungal infection (Zhao et al., 2013). We used the dbCAN2 pipeline to annotate CAZymes in the genome of strain HN2 (Drula et al., 2022). We annotated 594 CAZymes in the genome, including 107 auxiliary activity family proteins, 268 glycoside hydrolases,104 glycosyltransferases, 23 polysaccharide lyases, 43 carbohydrate esterases, and 15 carbohydrate-binding module family proteins. In addition, 34 genes have dual functions and are classified into two categories of CAZymes.

**Fig. 1.**
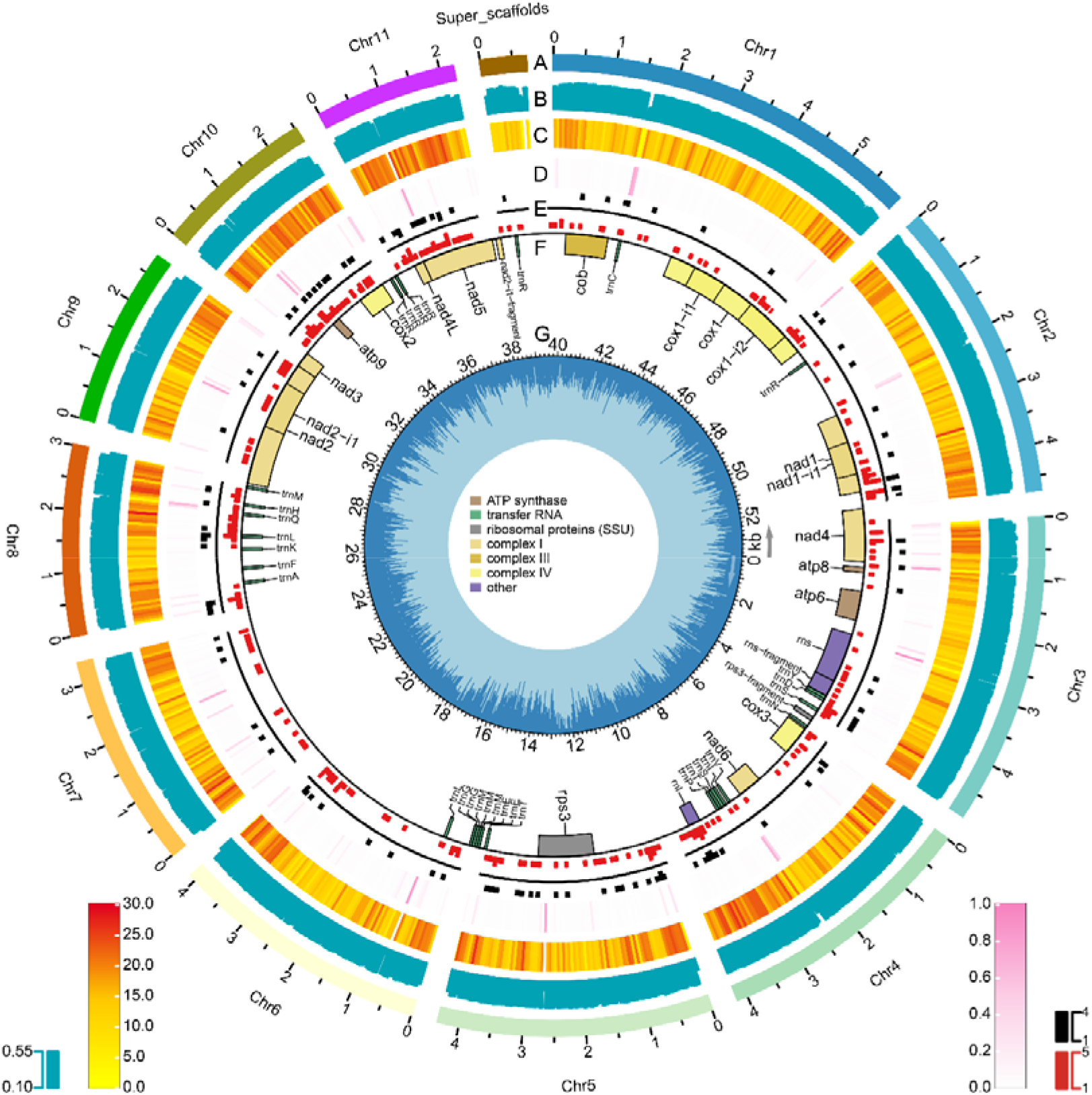
Features of the nuclear and mitochondrial genomes of *F. verticillioides* strain HN2. A. The 11 chromosomes and the supernumerary scaffold (super_scaffolds). B. GC content. C. Gene count. D. Repeat density. E. Candidate effectors (red) and conserved candidate effectors (black). Data in circles are displayed in nonoverlapping 50-kb intervals. F. Genes of the mitochondrial genome. ATP synthases (brown), transfer RNAs (green), ribosomal proteins (grey), oxidative phosphorylation (yellow), and others (purple). G. GC content of the mitochondrial genome.

Using GetOrganelle (v1.7.6.1) (Jin et al., 2020), we assembled the 53,764 bp complete mitochondrial genome of *F. verticillioides* strain HN2, with GC content at 33%. Codon preference analyses showed that codon TAT, ATA, and TTA are the most frequently used (Supplementary Table S6) (Rice et al., 2000). A total of 49 mitochondrial genes were annotated using GeSeq (Tillich et al., 2017), including 15 protein-coding genes, 32 tRNAs, and 2 rRNAs (*rns* and *rnl*), similar to the published results of *F. verticillioides* strain 7600 (Al-Reedy et al., 2012). A total of 52 simple sequence repeats (SSRs) were identified in the mitochondrial genome of *F. verticillioides* strain HN2 (Beier et al., 2017). The mitochondrial genome was visualized using software Chloroplot (Zheng et al., 2020).

In summary, we report the gap-free nuclear and mitochondrial genomes of *F. verticillioides* strain HN2, which will serve as an important resource for studies of *F. verticillioides*-maize interactions, and contribute to effective control of FER and FSR.

## Data availability

The HN2 genome file is available at NCBI under the project accession number PRJNA866387. The genome annotation file, the sequence file of candidate effectors, the mitochondrial genome file, the mitochondrial genome annotation file, the one-to-one orthologs list, the Nanopore and next-generation sequencing data have been deposited to Figshare database (https://doi.org/10.6084/m9.figshare.20431578).

## Author-Recommended Internet Resources

NextDenovo: https://github.com/Nextomics/NextDenovo

RepeatMasker: http://www.repeatmasker.org

Funannotate: https://github.com/nextgenusfs/funannotate

antiSMASH: https://antismash.secondarymetabolites.org/#!/about

GeSeq: https://chlorobox.mpimp-golm.mpg.de/geseq.html

