## Supplemental Files for "Gap-free nuclear and mitochondrial genomes of *Fusarium verticillioides* strain HN2"

**Supporting Information**

**Supporting Figures**

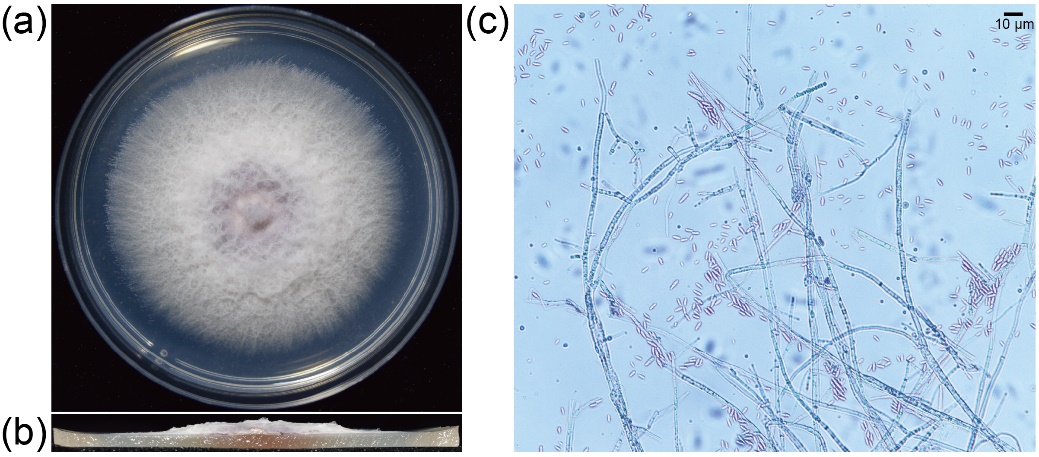

**Figure S1** **Morphological features of *F. verticillioides* strain HN2.**

(a) Five-day-old culture of strain HN2 on the potato dextrose agar (PDA) plates under ambient light, at 28°C. (b) Cross-section of the colony shown in (a). (c) Mycelia and microconidia of strain HN2.

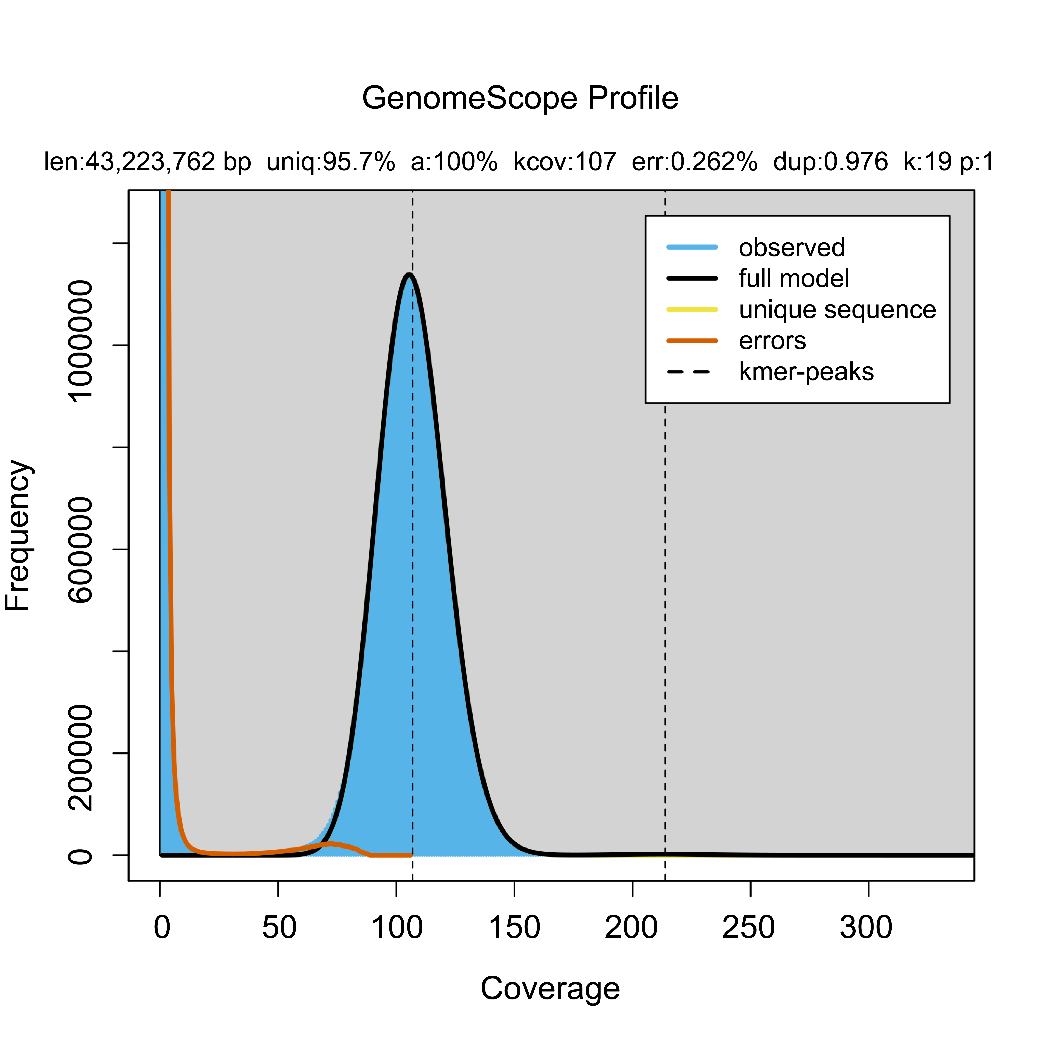

**Figure S2 The *k*-mer analysis of the genome of *F. verticillioides* strain HN2.**

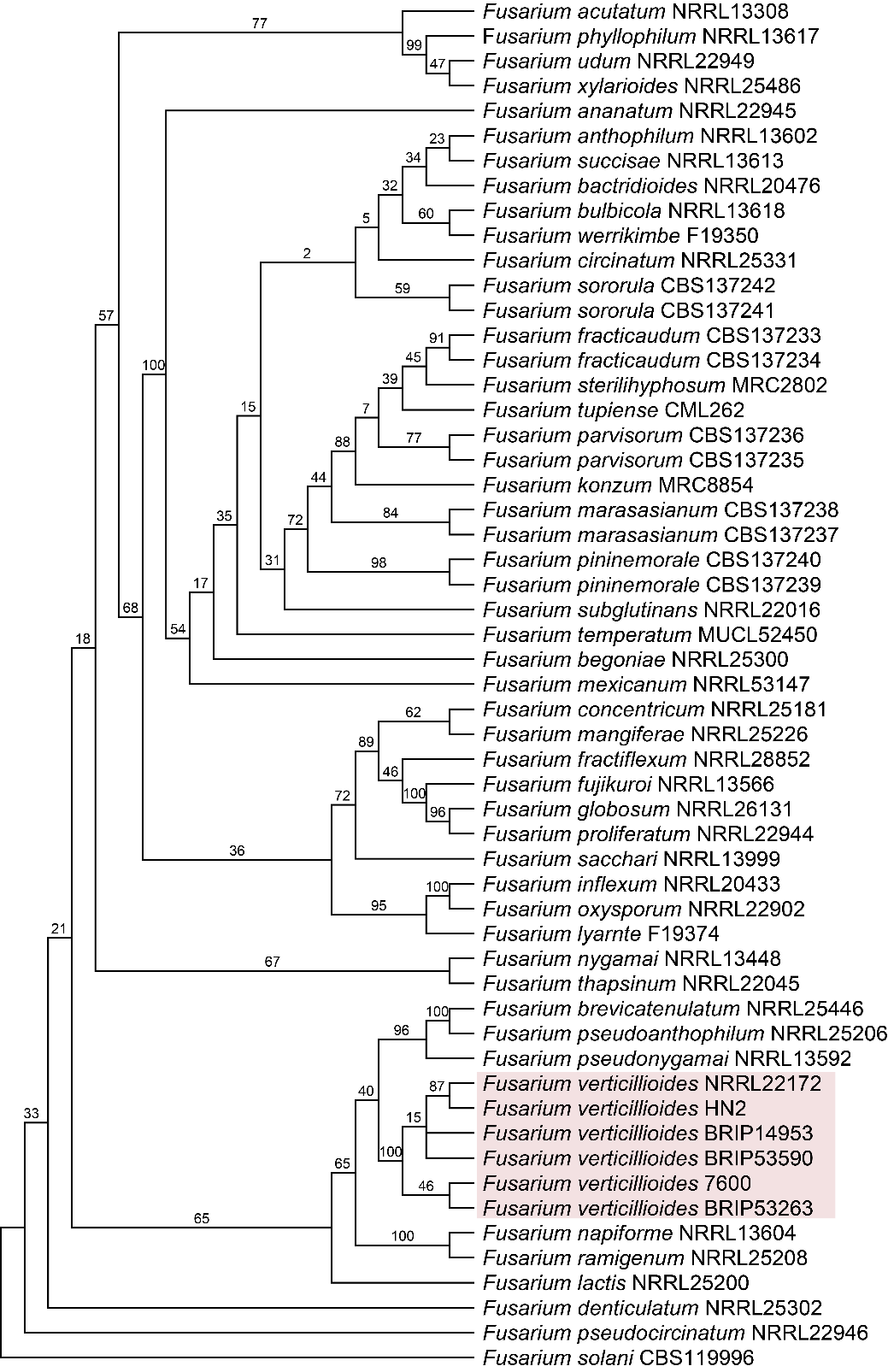

**Figure S3 Phylogenetic tree derived by maximum parsimony analysis.**

The phylogenetic tree was constructed based on translation elongation factor 1-α and β-tubulin genes. Strain HN2 was identified as *F. verticillioides* within the *Fusarium fujikuroi* species complex, highlighted in pink. *Fusarium* *solani* was used as the outgroup.

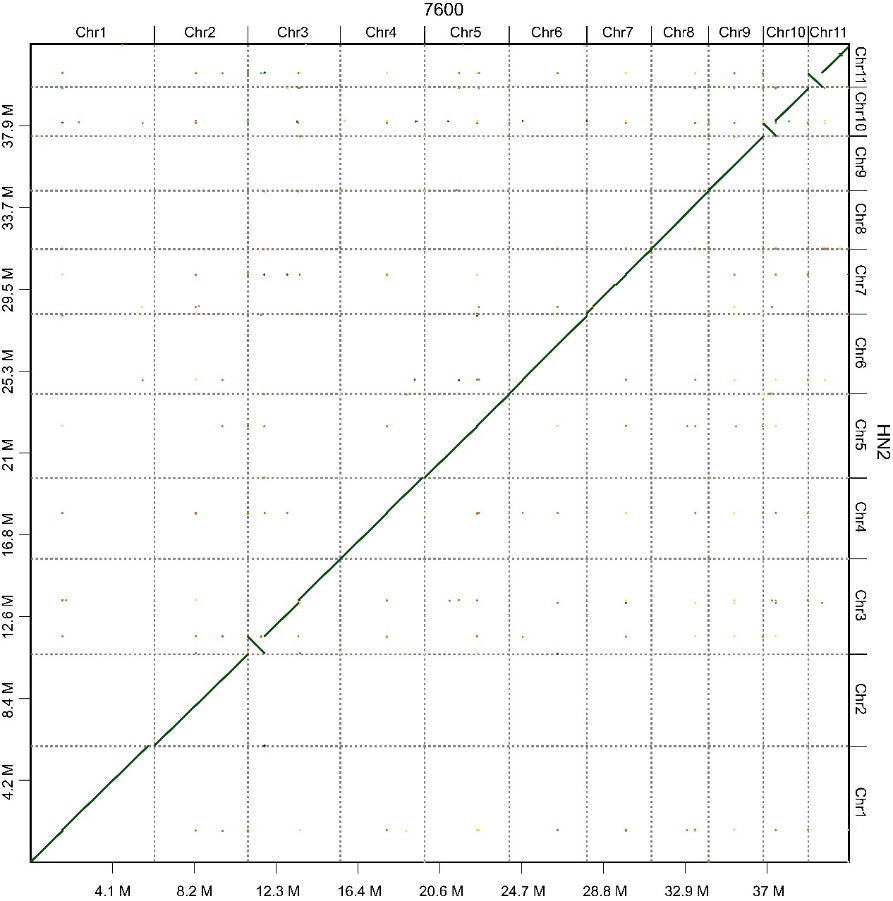

**Figure S4 Dot plot comparing the genomes of *F. verticillioides* strains HN2 and 7600.**

**Supporting Tables**

**Table S1 Telomere sequences in the genome of *F. verticillioides* strain HN2.**

| Chr | Chr-length | Start | End | Length | Type |
| --- | --- | --- | --- | --- | --- |
| Chr1 | 5,972,143 | 5,972,070 | 5,972,141 | 72 | TTAGGG |
| Chr2 | 4,724,490 | 4,724,369 | 4,724,476 | 108 | TTAGGG |
| Chr3 | 4,905,821 | 19 | 84 | 66 | CCCTAA |
| Chr4 | 4,158,290 | 4,158,252 | 4,158,275 | 24 | TTAGGG |
| Chr6 | 4,101,700 | 14 | 133 | 120 | CCCTAA |
| Chr6 | 4,101,700 | 4,085,522 | 4,085,587 | 66 | TTAGGG |
| Chr7 | 3,346,292 | 23 | 100 | 78 | CCCTAA |
| Chr7 | 3,346,292 | 3,346,198 | 3,346,287 | 90 | TTAGGG |
| Chr9 | 2,810,142 | 14 | 97 | 84 | CCCTAA |
| Chr10 | 2,518,577 | 15 | 122 | 108 | CCCTAA |
| Chr11 | 2,215,594 | 13 | 126 | 114 | CCCTAA |
| Chr11 | 2,215,594 | 2,215,524 | 2,215,589 | 66 | TTAGGG |
| Super_scaffolds | 721,913 | 5 | 58 | 54 | CCCTAA |
| Super_scaffolds | 721,913 | 65 | 112 | 48 | CCCTAA |
| Super_scaffolds | 721,913 | 721,810 | 721,911 | 102 | TTAGGG |

**Table S2 Repeat elements in the genome of *F. verticillioides* strain HN2.**

| Type | Number | Percentage of genome (%) |
| --- | --- | --- |
| LTR elements | 255 | 0.93 |
| DNA element | 505 | 0.81 |
| Unclassified | 2,118 | 1.00 |
| Small RNA | 123 | 0.09 |
| Simple repeats | 6,825 | 0.64 |
| Low complexity | 776 | 0.09 |
| Total content | 10,602 | 3.56 |

**Table S3 BUSCO assessment of the genomes of *F. verticillioides* strains HN2 and 7600**.

| Type | HN2-number | 7600-number |
| --- | --- | --- |
| Complete BUSCOs (C) | 752 (99.2%) | 747 (98.5%) |
| Complete and single-copy BUSCOs (S) | 750 (98.9%) | 746 (98.4%) |
| Complete and duplicated BUSCOs (D) | 2 (0.3%) | 1 (0.1%) |
| Fragmented BUSCOs (F) | 2 (0.3%) | 5 (0.7%) |
| Missing BUSCOs (M) | 4 (0.5%) | 6 (0.8%) |
| Total BUSCO groups searched | 758 | 758 |

**Table S4 Distribution of SNPs/InDels/SVs between strains HN2 and 7600.**

| Type | SNPs | InDels | SVs |
| --- | --- | --- | --- |
| Downstream | 16,069 | 2,059 | 846 |
| Exonic | 105,459 | 5,077 | 435 |
| Exonic; Splicing | 16 | 0 | 0 |
| Intergenic | 11,958 | 1,291 | 948 |
| Intronic | 13,356 | 2,188 | 61 |
| NcRNA_exonic | 92 | 27 | 8 |
| NcRNA_UTR5 | 0 | 1 | 0 |
| Splicing | 192 | 57 | 7 |
| Upstream | 25,235 | 3,481 | 764 |
| Upstream; Downstream | 46,075 | 6,236 | 566 |
| UTR3 | 12,074 | 1,851 | 100 |
| UTR5 | 10,594 | 2,250 | 55 |
| UTR5; UTR3 | 314 | 59 | 3 |
| SUM | 241,434 | 24,577 | 3,793 |

**Table S5 Major fungal pathogens of maize used for conservativeness analysis of effector proteins.**

| Pathogen | Strain | BioProject | Blast result |
| --- | --- | --- | --- |
| *Fusarium verticillioides* | 7600 | PRJNA15553 | 332 |
|  | BRIP14953 | PRJNA437508 | 332 |
| *Fusarium graminearum* | PH-1 | PRJEB5475 | 272 |
| *Bipolaris maydis* | C5 | PRJNA42739 | 134 |
| *Exserohilum turcicum* | Et28A | PRJNA245152 | 131 |

**Table S6 Codon usage in the mitogenome of *F. verticillioides* strain HN2.**

| **Codon** | **AA** | **Fraction** | **Frequency** | **Number** |
| --- | --- | --- | --- | --- |
| TAT | Y | 0.778 | 49.997 | 896 |
| ATA | I | 0.526 | 47.821 | 857 |
| TTA | L | 0.377 | 45.477 | 815 |
| AAA | K | 0.66 | 45.198 | 810 |
| TTT | F | 0.763 | 42.576 | 763 |
| AAT | N | 0.788 | 37.833 | 678 |
| TAA | * | 0.495 | 36.661 | 657 |
| ATT | I | 0.365 | 33.201 | 595 |
| TAG | * | 0.373 | 27.677 | 496 |
| GAA | E | 0.61 | 24.608 | 441 |
| GCT | A | 0.505 | 24.162 | 433 |
| CTA | L | 0.197 | 23.715 | 425 |
| AAG | K | 0.34 | 23.325 | 418 |
| CTT | L | 0.189 | 22.711 | 407 |
| AGC | S | 0.263 | 20.646 | 370 |
| GTA | V | 0.393 | 19.642 | 352 |
| AGT | S | 0.248 | 19.474 | 349 |
| GAT | D | 0.757 | 19.307 | 346 |
| ATG | M | 1 | 16.405 | 294 |
| AGA | R | 0.345 | 16.405 | 294 |
| GTT | V | 0.327 | 16.35 | 293 |
| GAG | E | 0.39 | 15.736 | 282 |
| TCT | S | 0.198 | 15.568 | 279 |
| CCT | P | 0.482 | 14.676 | 263 |
| CAA | Q | 0.589 | 14.62 | 262 |
| TAC | Y | 0.222 | 14.285 | 256 |
| TTC | F | 0.237 | 13.225 | 237 |
| TTG | L | 0.11 | 13.225 | 237 |
| ACT | T | 0.372 | 13.002 | 233 |
| CAT | H | 0.669 | 12.499 | 224 |
| ACA | T | 0.338 | 11.83 | 212 |
| AGG | R | 0.248 | 11.774 | 211 |
| GGA | G | 0.311 | 11.104 | 199 |
| GCA | A | 0.225 | 10.769 | 193 |
| TGT | C | 0.536 | 10.435 | 187 |
| TCA | S | 0.131 | 10.267 | 184 |
| CAG | Q | 0.411 | 10.211 | 183 |
| AAC | N | 0.212 | 10.156 | 182 |
| ATC | I | 0.108 | 9.821 | 176 |
| GGT | G | 0.273 | 9.765 | 175 |
| TGA | * | 0.132 | 9.765 | 175 |
| GTG | V | 0.194 | 9.709 | 174 |
| TGC | C | 0.464 | 9.04 | 162 |
| GGG | G | 0.247 | 8.816 | 158 |
| CTG | L | 0.071 | 8.537 | 153 |
| TGG | W | 1 | 7.645 | 137 |
| CCA | P | 0.244 | 7.421 | 133 |
| CTC | L | 0.057 | 6.808 | 122 |
| CGA | R | 0.14 | 6.64 | 119 |
| GCC | A | 0.136 | 6.529 | 117 |
| TCG | S | 0.083 | 6.529 | 117 |
| GCG | A | 0.134 | 6.417 | 115 |
| GAC | D | 0.243 | 6.194 | 111 |
| CAC | H | 0.331 | 6.194 | 111 |
| TCC | S | 0.077 | 6.082 | 109 |
| GGC | G | 0.169 | 6.026 | 108 |
| ACC | T | 0.15 | 5.245 | 94 |
| ACG | T | 0.14 | 4.91 | 88 |
| CCC | P | 0.152 | 4.631 | 83 |
| CGT | R | 0.098 | 4.631 | 83 |
| CGC | R | 0.094 | 4.464 | 80 |
| GTC | V | 0.086 | 4.297 | 77 |
| CCG | P | 0.123 | 3.739 | 67 |
| CGG | R | 0.075 | 3.571 | 64 |
